# The evolution of fluoroquinolone resistance in *Salmonella* under exposure to sub-inhibitory concentration of enrofloxacin

**DOI:** 10.1101/2021.08.18.456923

**Authors:** Yufeng Gu, Lulu Huang, Cuirong Wu, Junhong Huang, Haihong Hao, Zonghui Yuan, Guyue Cheng

## Abstract

The evolution of resistance in *Salmonella* to fluoroquinolones (FQs) under a broad range of sub-inhibitory concentrations (sub-MICs) has not been systematically studied. This study investigated the mechanism of resistance development in *Salmonella enterica* serovar Enteritidis (*S*. Enteritidis) under sub-MICs of 1/128×MIC to 1/2×MIC of enrofloxacin (ENR), a widely used veterinary FQ. It was shown that the resistance rate and resistance level of *S*. Enteritidis varied with the increase of ENR concentration and duration of selection. qRT-PCR results demonstrated that the expression of outer membrane porin (OMP) genes, *ompF, ompC* and *ompD*, were down-regulated first to rapidly adapt and develop resistance of ≤ 4×MIC, and as the resistance level increased (≥8×MIC), the up-regulated expression of efflux pump genes, *acrB, emrB* amd *mdfA*, along with mutations in quinolone resistance-determining region (QRDR) gradually played a decisive role. Cytohubba analysis based on transcriptomic profiles demonstrated that *purB, purC, purD, purF, purH, purK, purL, purM, purN* and *purT* were the hub genes for the FQs resistance. ‘de novo’ IMP biosynthetic process, purine ribonucleoside monophosphate biosynthetic process and purine ribonucleotide biosynthetic process were the top three biological processes screened by MCODE. This study first described the dynamics of FQ resistance evolution in *Salmonella* under a long-term selection of sub-MICs of ENR *in vitro*. In addition, this work offers greater insight into the transcriptome changes of *S*. Enteritidis under the selection of ENR and provides a framework for FQs resistance of *Salmonella* for further studies.

## 1 Introduction

*Salmonella enterica* serovar Enteritidis (*S*. Enteritidis), a zoonotic foodborne pathogen, has been widely recognized as one of the most common causes of gastroenteritis in humans^[1]^. According to the report of World Health Organization, *S*. Typhimurium and *S*. Enteritidis are the most frequently isolated *Salmonella* serotypes from countries involved in the Global Foodborne Infections Network^[2]^. Fluoroquinolones (FQs) have been broadly applied in clinical practice for treating salmonellosis in both humans and animals^[3, 4]^. The emergence of resistance to FQs has become a critical problem in clinical treatment of salmonellosis^[5]^.

The mechanisms of FQs resistance in *Salmonella* include point mutations in quinolone resistant determining regions (QRDRs) in *gyrA, gyrB, parC* and *parE*^[6]^. Besides, decreased intake as well as increased efflux of FQs adds to the resistant phenotype of *Salmonella*. For example, changes in outer membrane porins (OMPs) (e.g. OmpC, OmpD and OmpF)^[7]^and elevated expression of multidrug resistance (MDR) efflux pumps (e.g. AcrAB, AcrEF, EmrAB, MdfA and MdtK)^[8]^ of *Salmonella* has been demonstrated as resistance mechanism to FQs for both clinical resistant isolates and resistant clones *de novo* selected by increasing concentrations (above MIC) of FQs *in vitro*^[9]^. However, the time sequence of the emergence of these various resistance mechanisms and the correlation with the level of resistance and the pressure of different antibiotic concentration is unclear and remains to be studied in detail.

Antimicrobials at sub-inhibitory concentrations (sub-MICs) are commonly found in patients, livestocks and the environment, often at a wide concentration ranging from 1/4 to 1/230 of the MIC^[10, 11]^. However, previous understanding of the resistance evolution process is mostly based on mutants selected by incrementally increasing antibiotic concentrations within mutant selection windows (MSW)^[12, 13]^. It has been shown that *de novo* mutants can be selected at sub-MIC of antimicrobials associated with several secondary effects, such as inducing the SOS response, stimulating the production of reactive oxygen species, increasing the frequency of errors in protein synthesis, increasing the rates of recombination and horizontal gene transfer, etc^[14-18]^.

Recent work has shown that the resistance mechanisms induced by sub-MIC exposure may be different compared to selection with antibiotics concentration above MIC. In *S*. Enteritidis, high-level resistance was selected by sub-MICs of streptomycin through multiple small-effect resistance mutations, whereas specific target mutations were generated under selection with antibiotics concentration above MIC^[19]^. While many studies have investigated the resistance mechanism of bacteria under a short-term exposure to antibiotics ^[20, 21]^, less is known about the effects of long-term exposure to sub-MIC of antibiotics. When exploring the *de novo* high-level or clinical resistance to the antimicrobial agent, most of these reports are endpoint observations and seldom take into account the changes occurring during the resistance evolution process. A more comprehensive understanding of the resistance development trajectory could help overcome resistance emergence.

Here, we systematically explored the resistance evolution of *S*. Enteritidis during a long-term exposure to a wide range of sub-MICs (1/128×MIC to 1/2×MIC) of enrofloxacin (ENR) and compared the effect of several concentration of ENR on the origin of resistance, focusing on the resistance mechanism to ENR. The known resistance mechanisms associated with *de novo* antibiotic resistance were analyzed in this study, including QRDRs of *gyrA, gyrB, parC* and *parE* genes, expression levels of the OMPs and MDR efflux pump genes. Transcriptome profiles of *S*. Enteritidis mutants with MIC level of 32×MIC, 16×MIC and 8×MIC were compared to *S*. Enteritidis parental strain, giving an indication of the resistance evolution route and molecular mechanism of *S*. Enteritidis under exposure to ENR in a long-term. Therefore, the purpose of this study was to determine the role of different resistance mechanism under selection of sub-MICs of ENR during the resistance development term. Overall, our findings added to evidence that sub-MIC antibiotic exposure and long-term selection prime bacteria for reduced susceptibility and resistance evolution.

## 2 Materials and methods

### 2.1 Bacteria, drugs, and reagents

*S*. Enteritidis CICC21527 was purchased from (China Center of Industrial Culture Collection, CICC, China). ENR (purity of 94.2%) was bought from China Institute of Pharmaceutical and Biological Products Inspection (Beijing, China). Luria-Bertani broth (LB) and Tryptone soybean agar (TSA) was purchased from HOPEBIO (Tsingtao, China). Premix Taq was bought from Moralsbio (Wuhan, China), and Ex Taq^Tm^ DNA Polymerase and SYBR was bought from Vazyme Biotech (Nanjing, China). HiFiScript gDNA Removal RT MasterMix was from Cwbio (Beijing, China), and RNAprep pure Bacteria kit was from Majorbio (Shanghai, China). gDNA Removal RT MasterMix was bought from Cwbio (Beijing, China).

### 2.2 Antimicrobial susceptibility testing

The MICs of ENR for wild-type and mutants of *S*. Enteritidis CICC21527 were determined using the broth micro-dilution method, according to the guidelines of the Clinical and Laboratory Standards Institute (CLSI)^[22]^.

### 2.3 *In vitro* selection of mutants under sub-MICs of ENR

To select *de novo* generated mutants, *S*. Enteritidis CICC21527 was cultured and passaged respectively in LB medium containing ENR at concentrations lower than MIC values, including 0.031 μg/mL (1/2×MIC), 0.016 μg/mL (1/4×MIC), 0.008 μg/mL (1/8×MIC), 0.004 μg/mL (1/16×MIC), 0.002 μg/mL (1/32×MIC), 0.001 μg/mL (1/64×MIC) and 0.0005 μg/mL (1/128×MIC). The culturing, passaging and mutant screening methods were carried as previously described^[23,24]^. The MICs of the selected mutants were confirmed by antimicrobial susceptibility testing. The 2×MIC mutants selected by 1/2×MIC, 1/4×MIC, 1/8×MIC, 1/16×MIC, 1/32×MIC, 1/64×MIC and 1/128×MIC of ENR were named 2M (1/2M), 2M (1/4M), 2M (1/8M), 2M (1/16M), 2M (1/32M), 2M (1/64M) and 2M (1/128M), respectively. The 4×MIC to 32×MIC mutants induced by sub-MICs of ENR were also similarly named. The mutants were grouped as reduced susceptibility (MIC=0.125-0.5 μg/mL) and resistance (MIC≥1 μg/mL), according to CLSI guidelines^[25]^.

### 2.4 Sequence analysis of QRDR region in *gyrA, gyrB, parC*, and *parE* genes

Strains 2M (1/2M), 2M (1/8M), 2M (1/32M), 2M (1/128M), 4M (1/2M), 4M (1/8M), 4M (1/32M), 4M (1/128M), 8M (1/2M), 8M (1/8M), 8M (1/32M), 8M (1/128M), 16M (1/2M), 16M (1/8M), 16M (1/32M) and 32M (1/2M) were applied to the detection of the QRDR region in *gyrA, gyrB, parC*, and *parE*, according to Kim *et al*^[26]^. The PCR products were purified from agarose gels using a TIANgel Purification Kit (TianGen BioTech Co. Ltd, China), followed by nucleotide sequencing performed by Sangon Biotech (Shanghai) Co. Ltd, China. The sequencing results were compared with the genome sequence of *S*. Enteritidis CICC21527 (SRA Accession No. SRR14246558).

### 2.5 Examination of the expression levels of OMPs and MDR efflux pump transporters

The strains as described in section **2.4** were subjected to gene expression analysis of *ompC, ompD, ompF, acrB, acrF, emrB, mdfA, and mdtK*. Total RNA was harvested from 1 mL aliquots of culture using RNAprep pure Bacteria kit according to the manufacturer’s recommendation. DNA in total RNA was removed by treatment with HiFiScript gDNA Removal RT MasterMix and cDNA synthesis was performed using HiFiScript gDNA Removal cDNA Synthesis Kit according to the method described in the manufacturer. qRT-PCR amplification was conducted with an initial step of 5 min at 95°C, followed by 40 cycles of 10 s at 95°C, 30 s at the annealing temperature at 60°C. The *gapA* gene was used as an internal control for normalization, and the parental strains were used as references for their derived mutants. The 2^−ΔΔCT^ method was used for relative gene expression calculations. Each RNA sample was tested in triplicate and the primers used are listed in **Tab. S1**.

### 2.6 RNA sequencing and bioinformatic analysis

The total RNA of parental *S*. Enteritidis CICC21527, reduced susceptibility mutant 8M (1/128M), and resistant mutants 16M (1/8M) and 32M (1/2M) was processed as the reference described^[24]^. The samples were paired-end sequenced using an Illumina HiSeq™ 2000 system (Personalbio technology Co. Ltd, Nanjing, China). The reference genome for annotation was *S*. Enteritidis CICC21527 genome (SRA Accession No. SRR14246558). The sequencing data was submitted to the National Center for Biotechnology Information Sequence Read Archive (SRA) under Accession No. PRJNA700473.

To characterize the biological pathways associated with the co-DEGs of ENR resistance, co-DEGs were analysed in the ClueGO. The Retrieval of Interacting Genes database online tool (STRING; http://stringdb.org/) was used to analyse the PPI of DEGs, and those experimentally validated interactions with a combined score >0.4 were selected as significant. The screened networks were visualized by Cytoscape 3.8.0. The Cytohubba was used to check the hub genes and the MCODE was performed to establish PPI network modules, Degree cutoff = 2, Node score cutoff = 0.2, k-core = 2, Max. Depth =100 as selected.

## 3 Results

### 3.1 Resistance development of *S*. Enteritidis under exposure to sub-MICs of ENR *in vitro*

The MIC of ENR for the parental *S*. Enteritidis CICC21527 strain was determined to be 0.0625 μg/mL. When exposed to sub-MICs of ENR, a gradual increase in the size of reduced susceptibility subpopulations was appeared during 600 generations, while no decrease in susceptibility was observed in the absence of ENR (**Fig. 1**). With the increasing of ENR concentration, the mutants were enriched faster. Except 1/64×MIC and 1/128×MIC induction groups, all of the lineages had subpopulations with MIC value higher than 1 μg/mL (16×MIC) after 600 generations. 32×MIC resistant subpopulations could only be selected by 1/2×MIC concentration of ENR at 600 generations. This showed an association between the concentrations of ENR and resistance occurrence rates as well as resistance levels of the mutant subpopulation.

**Fig. 1.**
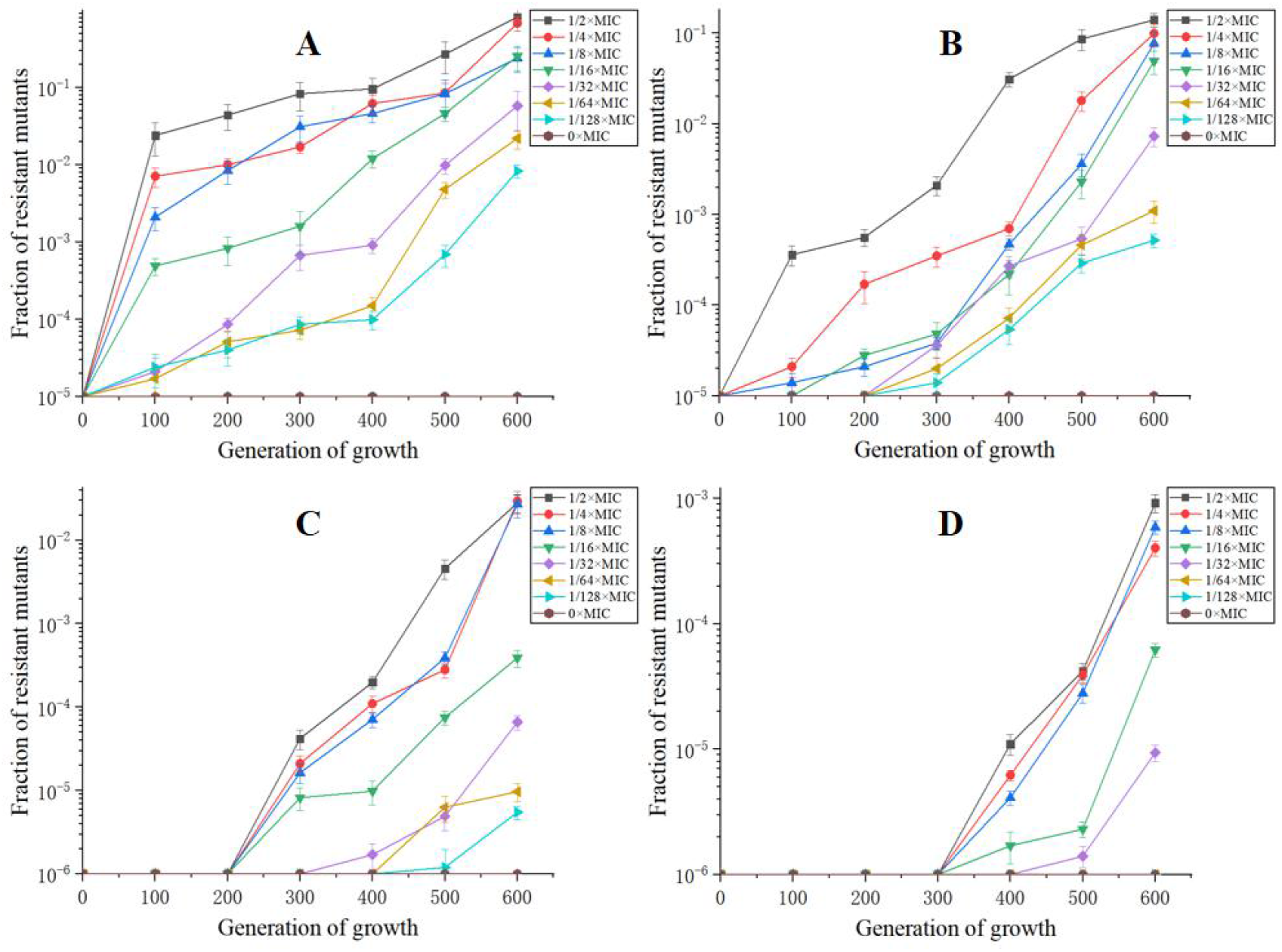
**Resistance rates of *S*. Enteritidis CICC21527 exposed to sub-MICs of ENR at resistant level of 2×MIC (A), 4×MIC (B), 8×MIC (C) and 16×MIC (D)**.

### 3.2 Mutations in the QRDRs of the mutants with reduced susceptibility to ENR

Compared to the parental strain, 12 out of 16 strains exhibiting MICs of 2 to 16×MIC had a mutation in the QRDR of the *gyrA* gene (**Tab. 1**). Among them, the mutation of Ser83Tyr was the most frequent (n=6), followed by the mutations of Ser83Phe (n=5) and Asp87Gly (n=2). It was also demonstrated that no mutation in *gyrA* were found in all reduced susceptibility mutants(≤8×MIC), while were found in all resistant mutants (≥ 16×MIC). No mutations in the QRDRs of *gyrB, parC* and *parE* were observed.

**Tab. 1.** Mutation sites in the QRDRs of *gyrA, gyrB, parC*, and *parE* genes of *S*. Enteritidis mutants.

| Strain NO. | MIC (μg/mL) | Substitutions in QRDR amino acid residues |  |  |  |
| --- | --- | --- | --- | --- | --- |
|  |  | *gyrA* | *gyrB* | *parC* | *parE* |
| 2M(1/2M) | 0.125 | wt | wt | wt | wt |
| 2M(1/8M) | 0.125 | Asp87Gly | wt | wt | wt |
| 2M(1/32M) | 0.125 | wt | wt | wt | wt |
| 2M(1/128M) | 0.125 | Ser83Tyr | wt | wt | wt |
| 4M(1/2M) | 0.25 | wt | wt | wt | wt |
| 4M(1/8M) | 0.25 | Ser83Phe | wt | wt | wt |
| 4M(1/32M) | 0.25 | Ser83Tyr | wt | wt | wt |
| 4M(1/128M) | 0.25 | Ser83Tyr | wt | wt | wt |
| 8M(1/2M) | 0.5 | wt | wt | wt | wt |
| 8M(1/8M) | 0.5 | Ser83Phe | wt | wt | wt |
| 8M(1/32M) | 0.5 | Ser83Phe | wt | wt | wt |
| 8M(1/128M) | 0.5 | Asp87Gly | wt | wt | wt |
| 16M(1/2M) | 1 | Ser83Phe | wt | wt | wt |
| 16M(1/8M) | 1 | Ser83Phe | wt | wt | wt |
| 16M(1/32M) | 1 | Ser83Tyr | wt | wt | wt |
| 32M(1/2M) | 2 | Ser83Tyr | wt | wt | wt |

Note: "wt" represented no mutation was observed.

### 3.3 Expression of OMPs and MDR efflux pump transporters of the mutants with reduced susceptibility to ENR

The expression of the OMP genes, *ompC, ompD, ompF* and genes encoding MDR efflux pump transporters, *acrB, acrF, emrB, mdfA, mdtK* of mutants was shown in **Fig. 2**. In the mutants with susceptibility level less than 8×MIC, the expression of *ompC, ompD* and *ompF* were down regulated, and the amount of down-regulation decreased with the increase of resistance level. When the susceptibility level was more than 8×MIC, the expression of *ompC* and *ompD* shifted to up-regulation, while the expression of *ompF* remained down-regulated. The result showed that the expression of *ompF* was well correlated with the selected concentration of ENR.

**Fig. 2.**
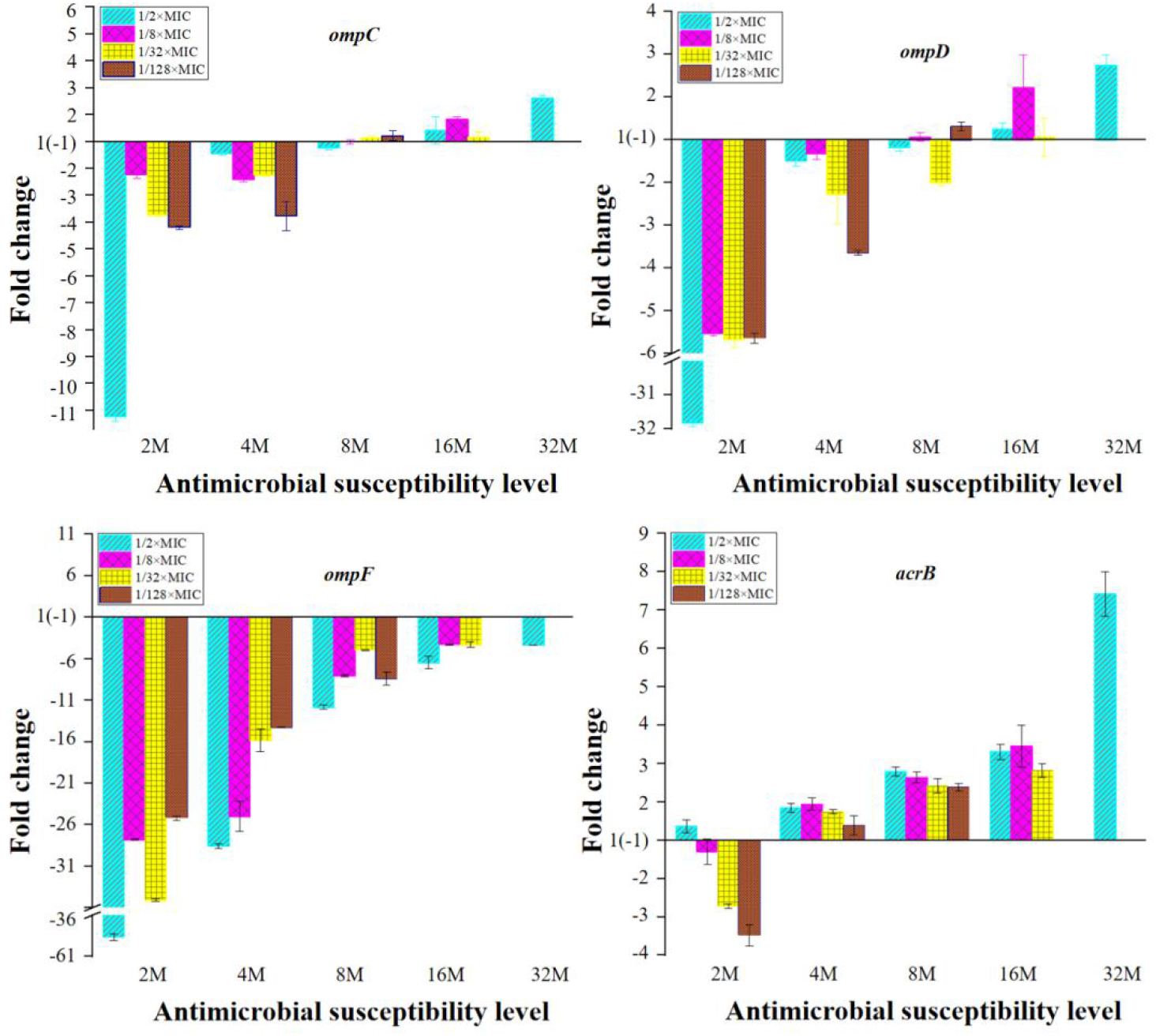
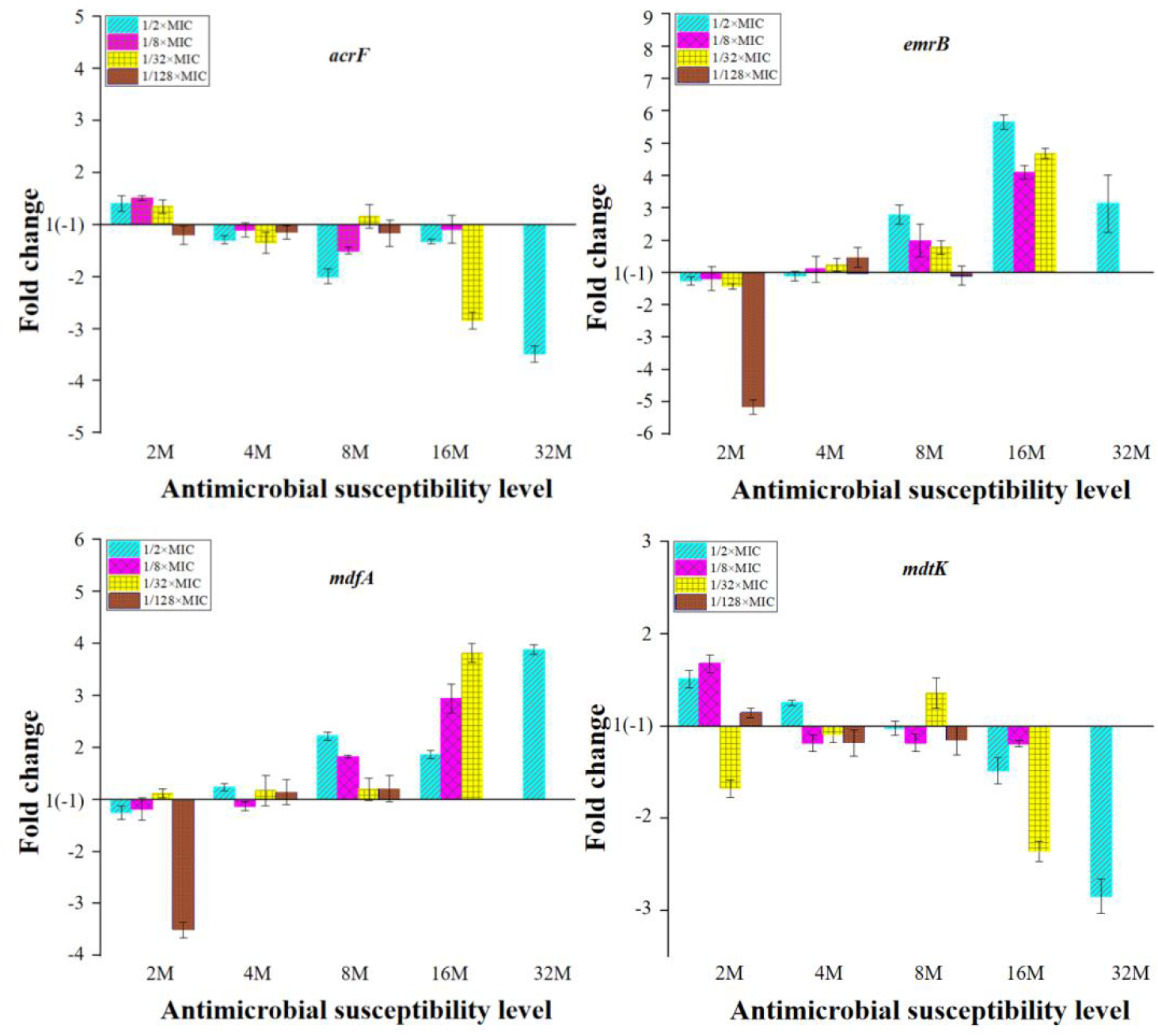
mRNA expression levels of the OMPs and MDR efflux pump transporter genes in *Salmonella*. Fold change = 2^−ΔΔCT^, ΔΔCt = (CT _target_ – CT _gapA_) mutant – (CT _target_ – CT _gapA_) parental. 2M, 4M, 8M, 16M represent 2×MIC mutants, 4×MIC mutants, 8×MIC mutants, 16×MIC mutants, respectively.

In general, *acrB, emrB* and *mdfA* were down regulated in the 2×MIC mutants. When the susceptibility level was equal or greater than 4×MIC, these three genes turned to up-regulated expression, and the expression level increased with the increase of resistant level with *acrB* gene exhibiting a higher level of up-regulation compared to those of the *emrB* and *mdfA* genes (**Fig. 2**). The expression of the other two MDR efflux pump transporter genes, *acrF* and *mdtK*, displayed a more strain dependent pattern in the reduced susceptible mutants, most of which showed up-regulation of *acrF* and *mdtK* genes in the 2×MIC mutants and down-regulation in mutants with resistant level ≥4×MIC, and as the resistant level increased, the expression of these two genes gradually decreased.

### 3.4 Transcriptomic profiles of *S*. Enteritidis mutants induced by sub-MICs of ENR

Reduced susceptible mutant 8M (1/128M) (Group E), resistant mutant 16M (1/32M) (Group D) and 32M (1/2M) (Group C), and parental strain (Group B) were selected for analysis of transcriptomic profiles. The pearson correlation coefficient in each group was greater than 0.91, indicating that the correlation between triplicate samples in the same group was good (**Fig. S1**). Compared to the parental strain, 2040 DEGs (1032 up-regulated and 1008 down-regulated) were found in the resistance mutant 32M (1/2M), 1497 DEGs (723 up-regulated and 774 down-regulated) in resistant mutant 16M (1/32M) and 1196 DEGs (644 up-regulated and 552 down-regulated) in reduced susceptibility mutant 8M (1/128M). Compared to the parental strain, there were 573 co-differentially expressed genes (co-DEGs) among the three mutants; 333 genes were up-regulated and 240 genes were down-regulated in mutant 32M (1/2M); 300 genes were up-regulated and 273 genes were down-regulated in mutant 16M (1/32M); 298 genes were up-regulated and 275 genes were down-regulated in mutant 8M (1/128M).

The 573 co-DEGs were enriched in 24 GO terms, including ribosome, purine nucleobase biosynthetic process, etc (**Fig. S2**). Ninety-six common KEGG Pathways were obtained, including ribosome, arginine and proline metabolism, nitrotoluene degradation, Lysine degradation, tryptophan metabolism, fructose and mannose metabolism, PTS system, etc (**Fig. S3**). Based on the information in the STRING protein query from public databases, 338 co-DEGs were mapped with the reference species of *S*. enterica CT18. Then 120 genes were obtained probably related to the mechanism of FQs resistance according to the annotation of Non-Redundant Protein Sequence Database. GO function (Kappa score ≥0.8) and KEGG pathway (P ≤0.05) enrichment analyses of 120 candidate co-DEGs were performed with clueGO (**Fig. 3)**. It was shown that these genes were classified into 14 functional categories including nucleoside metabolic, purine nucleobase biosynthetic process, nuclebase-containing compound biosynthetic process, hydroxymethyl-,formyl- and related transferase activity, tricarboxylic acid cycle, short-chain fatty acid metabolic, nuclebase-containing compound metabolic process, chromosome, DNA topological change, purine ribonucleoside triphosphate binding, organelle organization, RNA binding, RNA catabolic process, purine-containing compound biosynthetic process (**Fig. 3A**). The metabolic pathways were significantly enriched in one carbon pool by folate, purine metabolism, propanoate metabolism, citrate cycle (TCA cycle) and RNA degradation pathways (**Fig. 3B**).

**Fig. 3.**
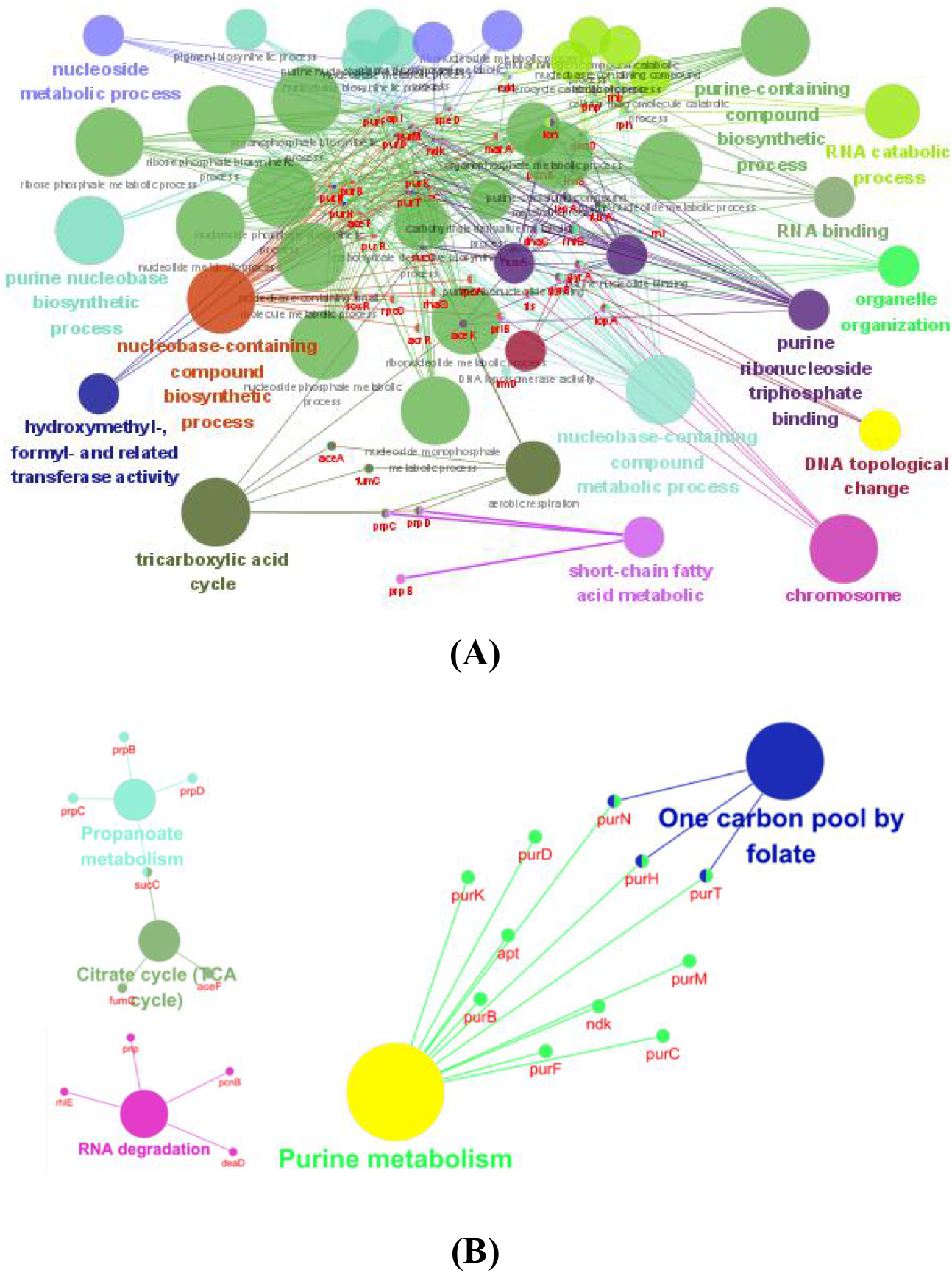
Enrichment analysis of GO and KEGG of genes that may be related to drug resistance by ClueGO. GO term enrichment (A); KEGG enrichment (B).

The 120 genes mentioned above encoding proteins belonging to the oxidoreductase, purine and pyrimidine metabolism, cell division, transcriptional regulator, stress response protein, DNA topoisomerase, DNA and RNA polymerase, RND efflux transporter were screened to identify molecular determinants associated with the response to ENR in *Salmonella*. With the aim of identifying key or central genes in the co-DEGs network of the *S*. Enteritidis mutants after exposure to sub-MICs of ENR, an analysis of hub gene identification was conducted based on STRING database (**Fig. 4A**).

**Fig. 4.**
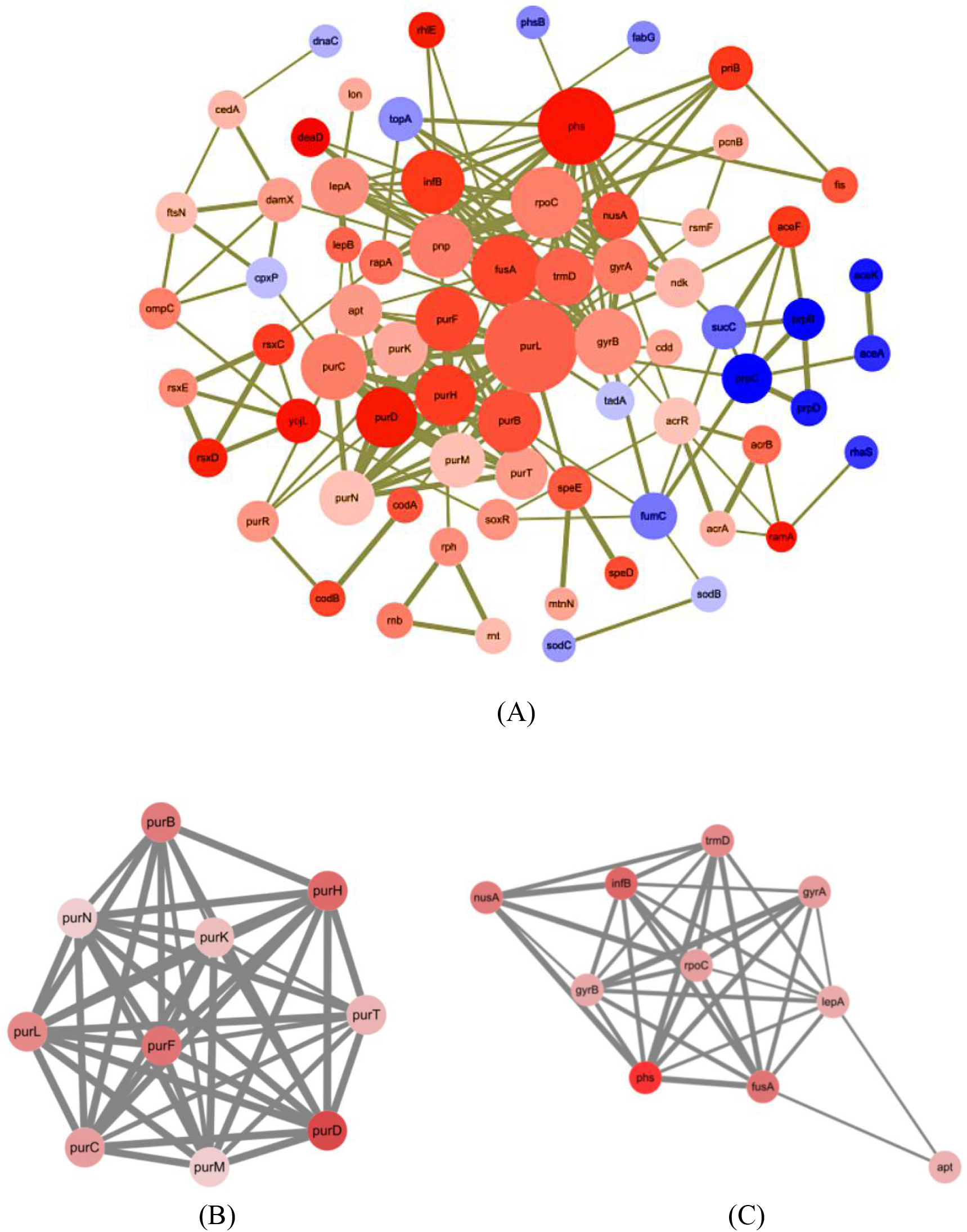
Network analysis of co-DEGs selected for underlying resistance mechanism. (A) Using the STRING online database, total of 120 co-DEGs were filtered into the PPI network and visualized by Cytoscape; (B, C) Top two PPI networks in MCODE analysis.

Furthermore, the Cytohubba result showed that *purB, purC, purD, purF, purH, purK, purL, purM, purN* and *purT* were the hube genes that responded to sub-MICs of ENR. To better understand the potential biological mechanism related to the network, screened the top two clusters was screened by MCODE with the highest clustering scores (Figure 4(B, C)) and the main biological processes (**Tab. 2**).

**Tab. 2.** Enrichment analysis of the top 2 MCODE genes function.

| MCODE | GO | Description | False discovery rate |
| --- | --- | --- | --- |
| MCODE-1 | GO:0006189 | 'de novo' IMP biosynthetic process | $4.08 \times 10^{-10}$ |
| MCODE-1/MCODE-2 | GO:0009168 | Purine ribonucleoside monophosphate biosynthetic process | $4.92 \times 10^{-9}$ |
| MCODE-1/MCODE-2 | GO:0009152 | Purine ribonucleotide biosynthetic process | $1.15 \times 10^{-8}$ |
| MCODE-1/MCODE-2 | GO:0034641 | Cellular nitrogen compound metabolic process | $1.55 \times 10^{-8}$ |
| MCODE-1/MCODE-2 | GO:0044271 | Cellular nitrogen compound biosynthetic process | $1.77 \times 10^{-8}$ |

The 39 DEGs out of the 120 co-DEGs selected by the criteria of expression fold-changes more than or equal to twice between these groups were selected for candidated key genes for their differential expression between mutants 32M (1/2M), 16M (1/32M) and 8M (1/128M) (**Tab. S1**). These genes were further screened and the heatmap was showed in **Fig. 5**, then STRING database was used to achieve the cluster map. Totally, ten clusters were identified including purine biosynthesis, purine biosynthesis, and pyrimidine metabolism, ‘de novo’ IMP biosynthetic process, response to antibiotic and transcription regulator, DNA topoisomerase, etc.

**Fig. 5.**
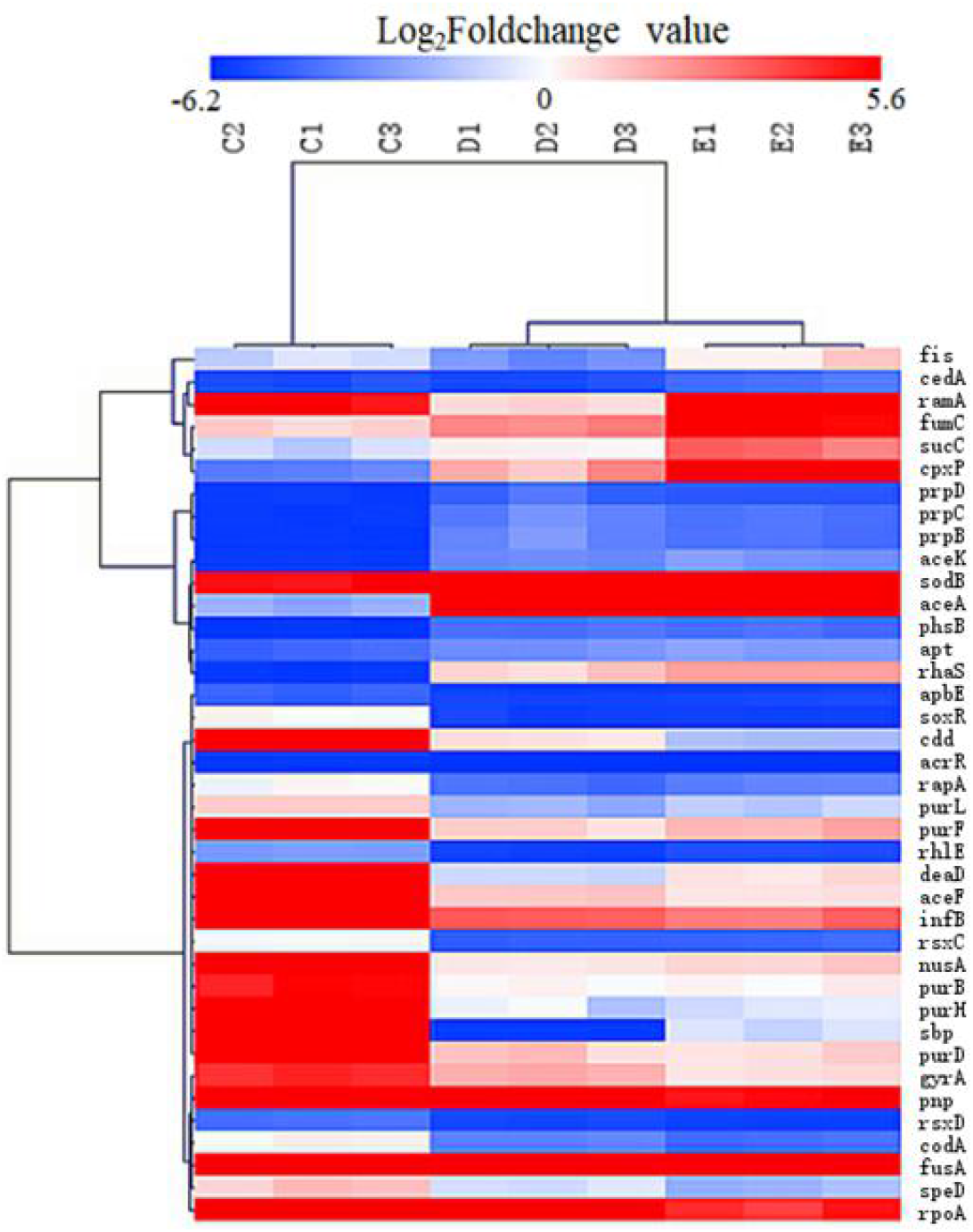
Heatmap of the candidated key genes involved in the sub-MIC induced ENR resistance in *S*. Enteritidis. “C_1_, C_2_, C_3_”, “D_1_, D_2_, D_3_”, “E_1_, E_2_, E_3_” represent triplicate of mutants 32M (1/2M),16M (1/32M) and 8M (1/128M) ; “red colour” represents gene up-regulation, “blue colour” represents gene down-regulation, and the shade of the color indicates the degree of gene expression.

The 573 co-DEGs were blasted in the CARD database, and the results showed that there were 19 known drug resistance genes (**Tab. 3**).

**Tab. 3.**
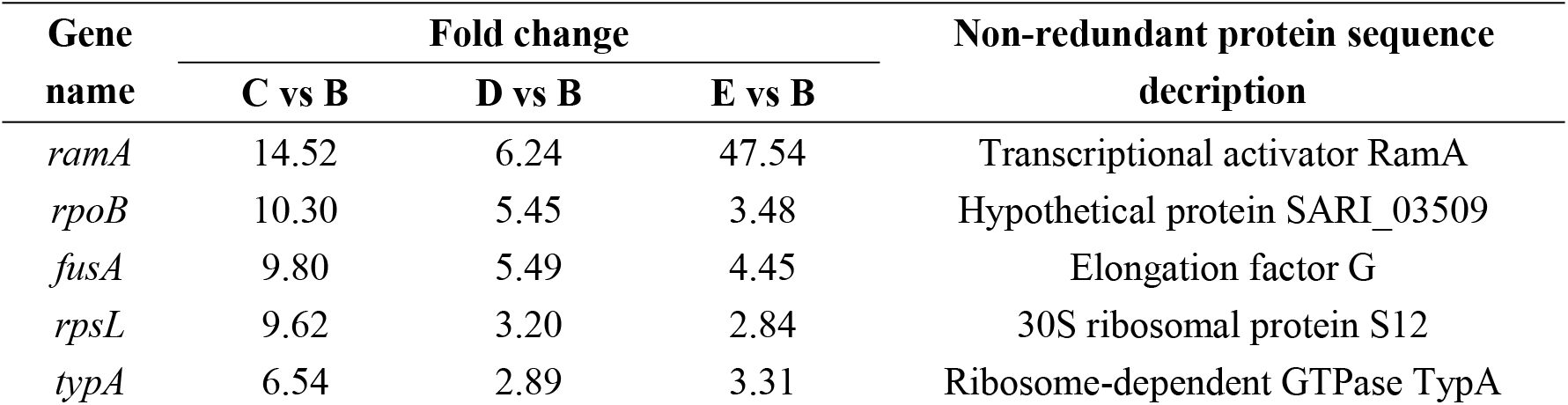
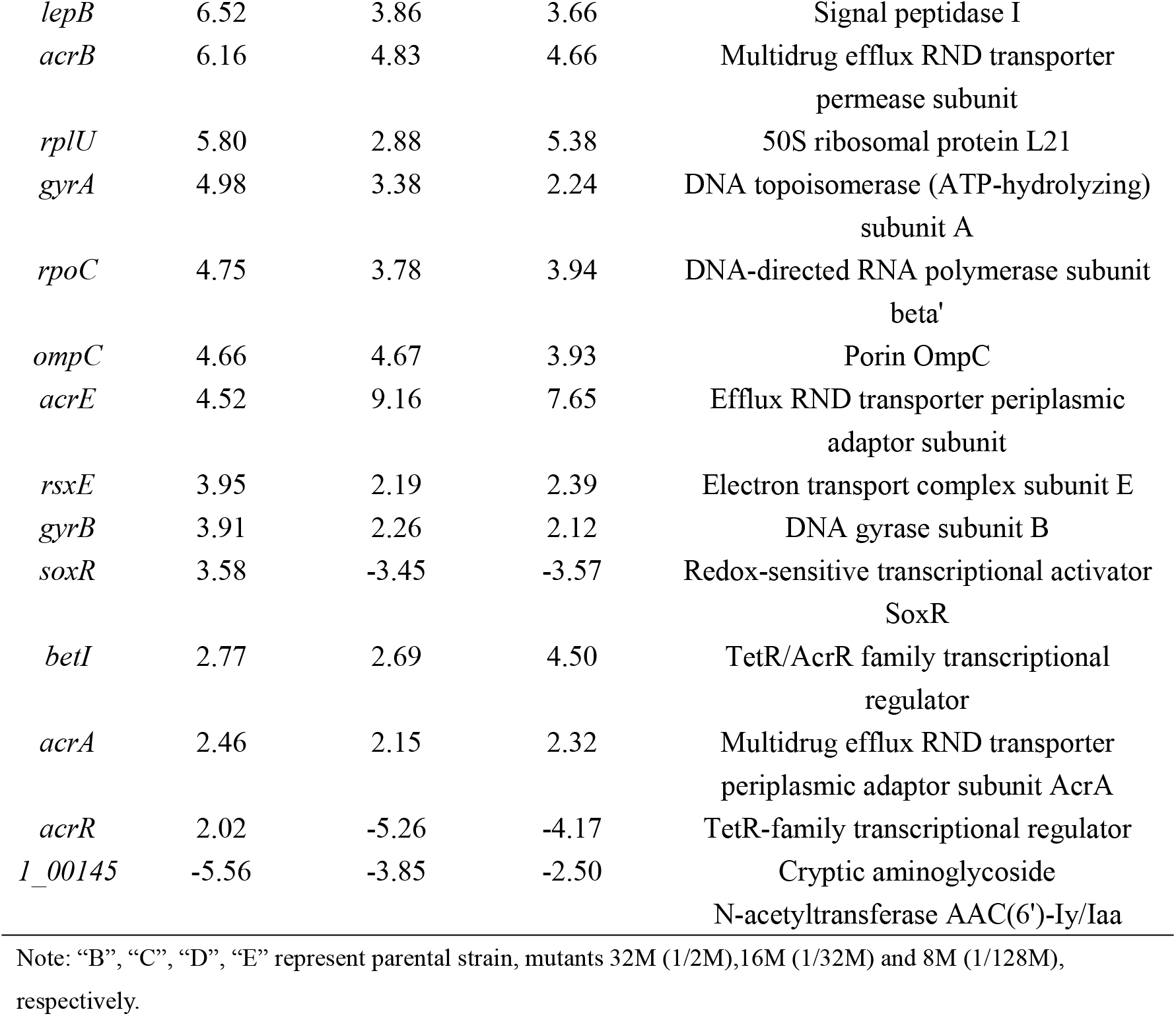
Resistance-related DEGs blasted in the CARD.

Based on the results of the transcriptomic analysis, the expression of the OMPs and MDR efflux pump transporter genes were presented in **Tab. 4**. The mRNA expression of OMPs (OmpA, OmpC, OmpD, and OmpF) was showed that only the *ompF* down-regulated in the reduced susceptibility mutant 8M (1/128M) and resistance mutant 32M (1/2M), so the decreasing OMPs permeability would not be a determining factors for mutants with resistance level ≥8MIC. Only *acrA, acrB, acrD, acrE, emrB, mdfA*, and *mdtB* genes had a significant up-regulation expression of MDR efflux pump genes compared with the parental strain. However, the MDR efflux pump genes of *acrF* and *mdtK* were not activated in mutants compared with the parental strain. The expression levels of *acrB* and *acrE* in mutants were much higher than other up-regulation genes; meanwhile, only the AcrAB efflux pump had two up-regulation subunits in all mutants (≥8MIC) compared with the parental strain. Our results indicated that overexpression of AcrAB efflux pump predominantly increase in resistance to ENR in mutants (≥8MIC), whereas AcrD, AcrEF, EmrAB, MdfA and MdtK efflux pump facilitated the reduced susceptibility to ENR in mutants (≥8MIC). These gene expression trends were generally consistent with the qRT-PCR results (**Fig. 2**).

**Tab. 4.** Expression levels of MDR efflux pump and OMPs in the transcriptome of *S*. Enteritidis mutants.

| Gene name | Fold Change |  |  | Non-redundant protein sequence description |
| --- | --- | --- | --- | --- |
|  | C vs B | D vs B | E vs B |  |
| *ompA* | 1.75 | 2.40 | 2.08 | Membrane protein |
| *ompC* | 4.66 | 4.67 | 3.93 | Porin OmpC |
| *ompD* | 1.17 | 15.41 | 6.80 | Porin OmpD |
| *ompF* | -2.38 | 1.06 | -3.45 | Porin OmpF |
| *mdsC* | -1.20 | -1.33 | -1.96 | Multidrug efflux transporter outer membrane subunit MdsC |
| *acrA* | 2.46 | 2.15 | 2.32 | Multidrug efflux RND transporter Periplasmic adaptor subunit AcrA |
| *acrB* | 6.16 | 4.83 | 4.66 | Multidrug efflux RND transporter permease subunit |
| *tolC* | 1.03 | 2.00 | 2.48 | Outer membrane protein TolC |
| *acrD* | 1.80 | 2.99 | 4.42 | Multidrug efflux RND transporter permease AcrD |
| *acrE* | 4.52 | 9.16 | 7.65 | Efflux RND transporter periplasmic adaptor subunit |
| *acrF* | -1.59 | -1.59 | -1.28 | Multidrug efflux RND transporter permease subunit |
| *emrA* | 1.22 | 1.34 | 1.12 | Multidrug efflux MFS transporter periplasmic adaptor subunit EmrA |
| *emrB* | 1.97 | 2.68 | 2.60 | Multidrug efflux MFS transporter permease subunit EmrB |
| *mdfA* | 1.33 | 1.88 | 2.98 | MFS transporter |
| *mdtK* | -1.49 | -1.19 | 1.35 | Multidrug efflux MATE transporter MdtK |
| *mdsA* | -1.79 | -2.22 | -2.08 | Multidrug efflux RND transporter periplasmic adaptor subunit MdsA |
| *mdsB* | -2.00 | -2.08 | -2.44 | Multidrug efflux RND transporter permease subunit MdsB |
| *mdtA* | 1.19 | -1.37 | 1.13 | Multidrug efflux RND transporter subunit MdtA |
| *mdtB* | 1.64 | 1.52 | 2.63 | Multidrug efflux RND transporter permease subunit MdtB |
| *mdtC* | -1.05 | 1.18 | 1.59 | Multidrug efflux RND transporter permease subunit MdtC |
| *macA* | -1.08 | 1.02 | 1.50 | Macrolide transporter subunit MacA |
| *macB* | 1.04 | 1.47 | 1.53 | Macrolide ABC transporter ATP-binding protein/permease MacB |

Note: The genes also detected in RT-PCR were shown in bold. “B”, “C”, “D”, “E” represent parental strain, mutants 32M (1/2M), 16M (1/32M) and 8M (1/128M), respectively.

## 4 Discussions

This study documented a versatile adaptive response of the *S*. Enteritidis under a long-term exposure to sub-MICs of ENR which resulted in a diversity of phenotypes including OMPs and MDR efflux pumps expression, QRDR mutation and transcriptomic changes. Mutations in the bacteria DNA gyrase (*gyrA* and *gyrB*) and topoisomerase IV (*parC* and *par*E) genes, as well as up-regulation of MDR efflux genes, were known to mediate bacterial resistance to FQs^[7, 26]^. In this study, the mutation of *gyrA* (Ser83Phe, Ser83Tyr, or Asp87Gly) was observed in all mutant strains except in reduced susceptible strains of 2M (1/2M), 2M (1/32M), 4M (1/2M) and 8M (1/2M) (**Tab. 1**). This is in consistence with the fact that the most common QRDR mutations occur in the *gyrA* gene, resulting in substitutions of Ser-83 with Tyr, Phe, or Ala, and of Asp-87 with Asn, Gly, or Tyr in *Salmonella* isolates^[26-28]^. Previous study demonstrated that point mutations were also observed in *parC* and *parE* with the concomitant presence of mutation in *gyrA* of *Salmonella* Paratyphi isolates with resistance to nalidixic acid^[29]^. It was also found that clinical *Salmonella* isolates evolved high-level of ciprofloxacin (CIP) resistance that was accompanied by additional mutations in GyrA and ParE^[30]^. Interestingly, no mutation was found in *gyrB, parC* or *parE* gene in our study, even in the higher level of resistance group (≥16MIC) (**Tab. 1**). One possible reason for this phenomenon was that the FQs resistance level of clinical isolates was much higher than the resistant level of the mutants which were selected in our study. Previous research showed that mutations in *gyrA* and *parC* genes confered a measurable fitness advantage over strains without these mutations^[31]^. According to the growth curve of *Salmonella* under exposure to a series of sub-MICs of ENR, it was revealed that the greater the selection pressure, the lower growth rates in our observation (**Fig. S4**). Because of the resistance level of resistants in this study was relatively low, so another reason might be that a single mutation in *gyrA* was sufficient to impose a loss of fitness. Several prior studies had shown that exposure to sub-MICs of CIP could select for first-step mutations that confer stable low-level resistance from both target mutations and efflux mechanism^[32, 33]^. In our study, no mutation was observed in *gyrA* gene selected by close-to-MIC concentrations (1/2×MIC) of ENR in all reduced susceptibility mutants in the early stage of resistance development (≤8MIC), but mutations of *gyrA* gene were obtained in all reduced susceptibility mutants except 2M(1/32M) selected by with low sub-MICs (≤1/4×MIC) of ENR (**Tab. 1**). This might occur because the mutants (≤8MIC) emerged fast under the close-to-MIC concentrations selection. In this process, no mutants had been selected in the population. So the initial adaptation manner of *Salmonella* included overexpression of efflux pumps and decrease of OMPs to rapidly emerge reduced susceptibility. While low sub-MIC of ENR had little influence on the survival of *Salmonella*, the effect of MDR efflux pumps was not obvious, and mutants with the susceptibility level of ≤ 8MIC were selected by a long-term. In addition, transcriptomic data showed that *gyrA* and *gyrB* were up-regulated in all mutants (**Tab. 3**). It was reported that the expression of *gyrA* and *parC* increased significantly in resistant *Salmonella enterica* serovar Typhimurium (*S*. Typhimurium) selected *in vivo*, but no changes in the expression of these genes were detected in *S*. Typhimurium selected *in vitro*^[12]^. Whether the up-regulated expression of these genes was a determinant of FQs resistance possibility required further investigation.

In addition, mechanisms affecting the cell envelope by increased/decreased expression of OMPs and/or efflux of FQs also contributed to the intracellular accumulation of FQs^[21, 34]^. In our study, The relative expression of outer membrane-related genes (*ompC, ompD* and *ompF*) were all down-regulated in the mutants with resistance level less than 8MIC, and the amount of down-regulation decreased with the increase of resistance level. Previous research showed that alterations in OMPs including disappearance of some or all of these proteins (OmpA, OmpC, OmpD and OmpF) enriched resistance to FQs in *Salmonella* isolates with the MIC value ≥32 μg/mL^[7]^. However, when the resistance level exceeds 8MIC, the *ompC* and *ompD* gene were overexpressed in all mutants, while the *ompF* gene was still suppressed in all mutants in our results (**Fig. 2**). This was also confirmed by the the transcriptomic results that the expression of porin-encoding genes (*ompA, ompC*, and *ompD*) except *ompF* were up-regulated in all mutants with susceptibility level ≥8MIC (**Tab. 4**). OmpF has been experimentally determined to be the most important porin in the resistant mutants selected by incrementally increasing CIP concentrations ^[35]^. In contrast to other antibiotics, ENR was reported to have higher affinities to OmpF channel in *Escherichia coli* (*E. coli*)^[36]^, and down-regulation of *ompF* had been associated with the decrease in the accumulation of FQs in *E. coli* ^[37, 38]^. Our data also showed that the down-regulation of *ompF* played the most important role in the initial stages of ENR resistance emergence.

It has been reported that the multidrug resistance (MDR) efflux pumps AcrAB-TolC, AcrEF, EmrAB, MdfABC and MdtK contributed to FQ resistance in *Salmonella*^[8]^. Our results revealed that AcrEF and MdtK efflux may have little contribution to ENR resistance at early stage, while AcrAB, EmrAB and MdfABC may play an important role in ENR resistance, since the expression level of *acrB, emrB, mdfA* was increased with increased level of FQs resistance and *acrB* gene was significantly increased, while the expression of the *acrF* and *mdtK* gene down-regulated as the susceptibility reduced (**Fig. 2**). This was also shown in the transcriptomic profiles that only *acrA, acrB, acrD, acrE, emrB, mdfA, mdtB* genes were significantly up-regulated (**Tab. 4**). Different performance of efflux pumps towards FQ pressure was also reported in the previous study that the expression level of *acrB* was increased and *acrF* decreased in CIP-resistants of *Salmonella* with the MIC value ≥2 μg/mL ^[34]^.

Previous study has shown that the *acrAB* or *acrEF* genes conferred multidrug resistance to numerous antibiotics, the *emrAB* gene conferred resistance to novobiocin and nalidixic acid, the *mdfA* gene conferred resistance to tetracycline, chloramphenicol, norfloxacin and doxorubicin and the *mdtK* gene conferred resistance to norfloxacin and doxorubicin in *S*. Typhimurium^[39]^. Therefore, we speculate that all these efflux pumps can efflux ENR, but there may be differences in substrate affinity between them, resulting in differences in their expression. Although MDR efflux pumps conferred only low-level resistance (2- to 8-fold increase in MIC values)^[40, 41]^, AcrB, EmrB, and MdfA were still working together with QRDR mutations beyond 16×MIC resistance levels. It was demonstrated in our results that as the expression of OMPs down-regulated, the expression level of *acrB, emrB, mdfA* were up-regulated, indicating OMP and MDR efflux pumps work alternately.

It was demonstrated that a feedback mechanism between nine homologous functional efflux pump genes through co-regulation of *ramA* and *marA* was found in *S*. Typhimurium^[42, 43]^. The marbox operon is responsible for producing the *marA, soxRS* and *ramA* transcriptional activator to activate *acrAB* transcription. But *acrR* is independent of *mar-sox-rob* for controlling the expression of *acrB* in *Salmonella*^[7, 44]^.

In our study, *ramA* were overexpressed in all mutants, while *soxR* and *acrR* gene were up-regulated in resistant mutant 32M (1/2M), but down-regulated in reduced susceptible mutant 8M (1/128M), resistant mutant 16M (1/32M) (**Tab. S2**). The overexpression of *marA* was only observed in resistant mutant 32M (1/2M), but difference expression in reduced susceptible mutant 8M (1/128M), resistant mutant 16M (1/32M). The expression of *ramA* was consistent with previous studies, and the differential expression of *soxR*, marA and *acrR* gene might be an important reason for the different expression levels of efflux pumps.

Beyond the role of target mutation, OMPs and MDR efflux pumps involved in FQs resistance, there is an increasingly recognized role for cellular processes such as purines metabolism. It was confirmed that purines metabolism are required for DNA and RNA synthesis^[45]^. Previous study showed that key genes involved in nucleotide biosynthesis were identified, including *purA* and *purD* in purine synthesis^[46]^. Another research showed that *purL* or *purM* mutant disrupted purine biosynthesis in *Burkholderia*^[47]^. It was also demonstrated that *purA* gene was up-regulated in olaquindox resistance *E. coli*^[24]^. Previous study showed that KEGG pathway of purine metabolism, pyrimidine metabolism was enriched in the proteomics analysis of FQs resistance *E. coli*^[48]^. The Cytohubba result showed that *purB, purC, purD, purF, purH, purK, purL, purM, purN* and *purT* were the hube genes and MCODE revealed that the main biological processes all involved in purine metabolism in this study (**Tab. 2**). This study have revealed that purine metabolism was the highly activate pathway by bioinformatics analysis. It remains to be determined whether purine metabolism and the other changes observed in the ENR mutant is a key pathway to FQs resistance.

## 5 Conclusions

In summary, this study shows an evolutionary process for salmonella on FQs resistance. mutants firstly decreased OMPs permeability to rapidly adapt the selected pressure circumstances in the initial stage of resistance emergence, then the expression of efflux pumps was up-regulated in the following process and QRDR mutation was obtained, resulting in a higher resistance level under a long-term selected pressure of the sub-MIC antibiotics *in vitro*. Hub genes (*purB, purC, purD, purF, purH, purK, purL, purM, purN* and *purT*) and the remarkable biological processes of purine metabolism were identified by bioinformatics analysis of transcriptomic profiles. This suggests that changes in FQs resistance based on gene expression patterns and metabolic pathways. However, the interplay between FQs resistance mechanisms and metabolic pathway requires further exploration.

## Abbreviations

CFU: Colony-Forming Units
CICC: China Center of Industrial Culture Collection
CIP: Ciprofloxacin
CLSI: Clinical and Laboratory Standards Institute
DEGs: Differentially expressed genes
*E. coli*: *Escherichia coli*
FQs: Fluoroquinolones
GO: Gene Ontology
KEGG: Kyoto Encyclopedia of Genes and Genomes
OMP: outer membrane porin
PCR: Polymerase Chain Reaction
LB: Luria-Bertani broth
MDR: multidrug resistance
co-DEGs: co-differentially expressed genes
QRDRs: quinolone resistant determining regions
*S*. Enteritidis: *Salmonella enterica serovar* Enteritidis
*S*. Typhimurium: *Salmonella enterica serovar* Typhimurium
ENR: Enrofloxacin
MIC: Minimum inhibitory concentration
Sub-MIC: Sub-inhibitory concentration
TSA: Tryptone soybean agar

## Acknowledgments

This work was supported by the National Natural Science Foundation of China (3207 2921 & 31502115).

## Author Contributions

Conceived and designed the experiments: YG GC. Performed the experiments: YG JH LH. Analyzed the data: YG HH GC. Contributed reagents/materials/analysis tools: GC ZY. Wrote the paper: YG LH CW GC.

## Supplementary data

**Figure S1.**
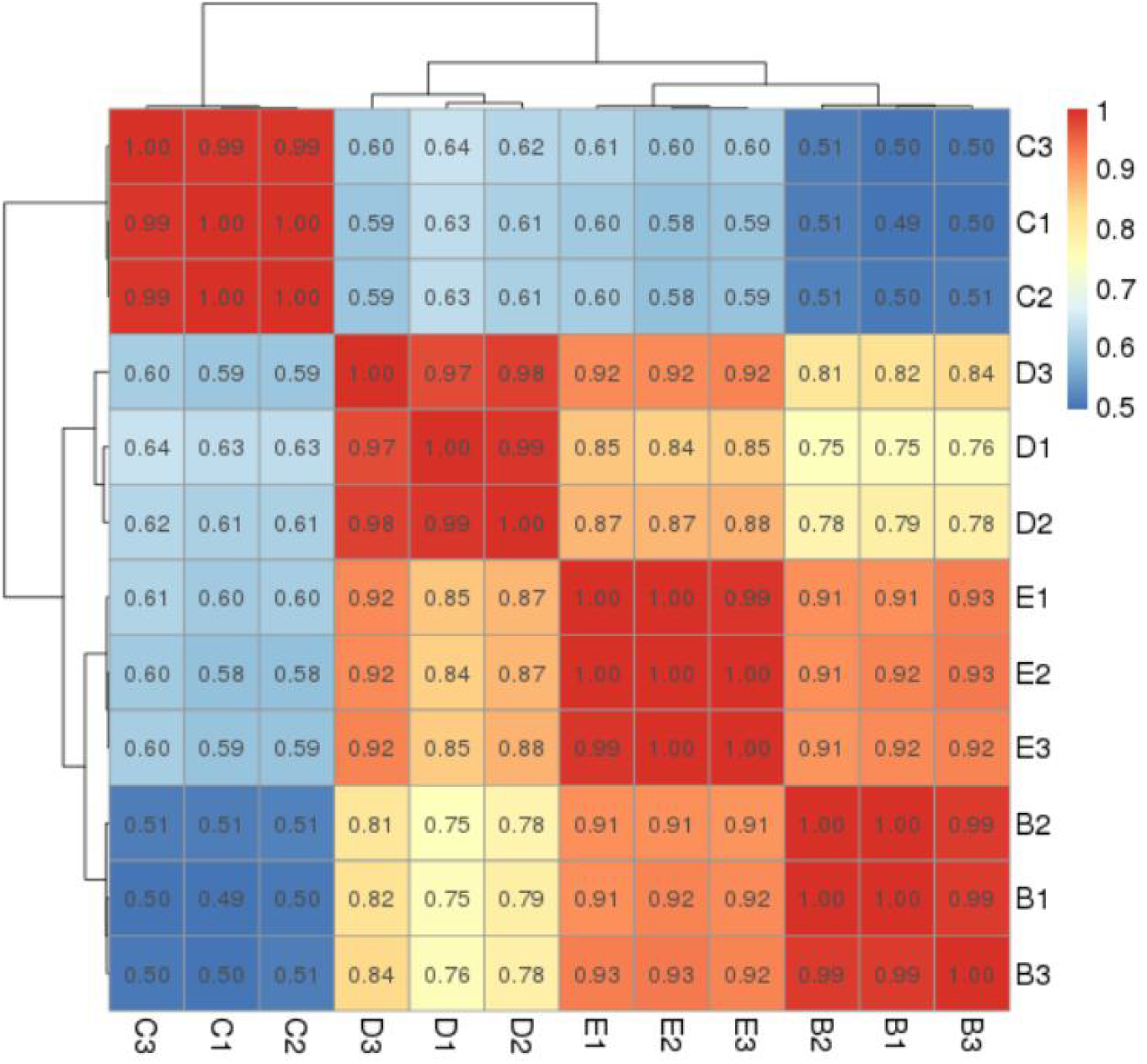
Correlation tests for the parental strain (Group B), strain 32M (1/2M) (Group C), strain 16M (1/8M) (Group D) and strain 8M (1/128M) (Group E) **(triplicates in each group)**. The abscissa and ordinate in the figure are sample numbers. The closer the block value is to 1, the higher the similarity is.

**Fig. S2.**
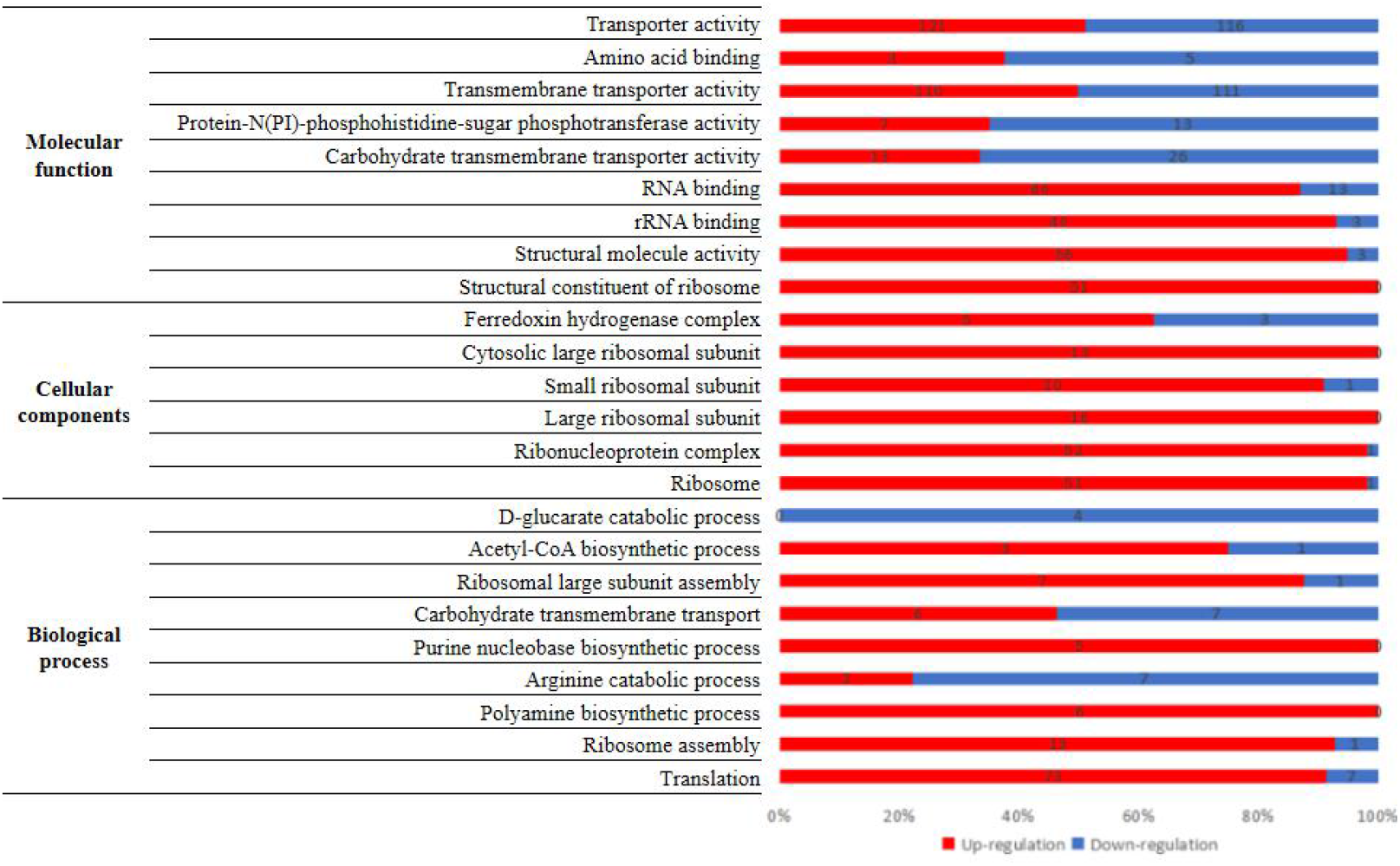
Enrichment of co-DEGs in GO Term.

**Fig. S3.**
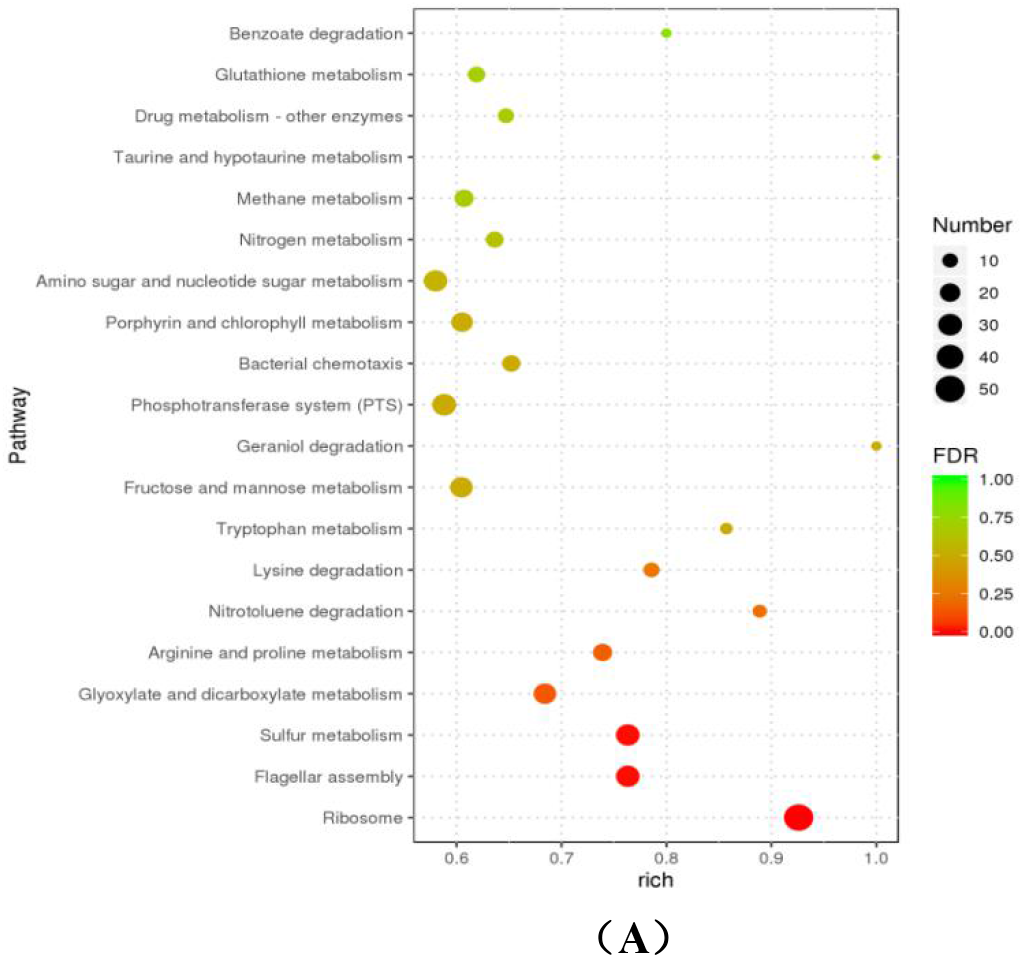
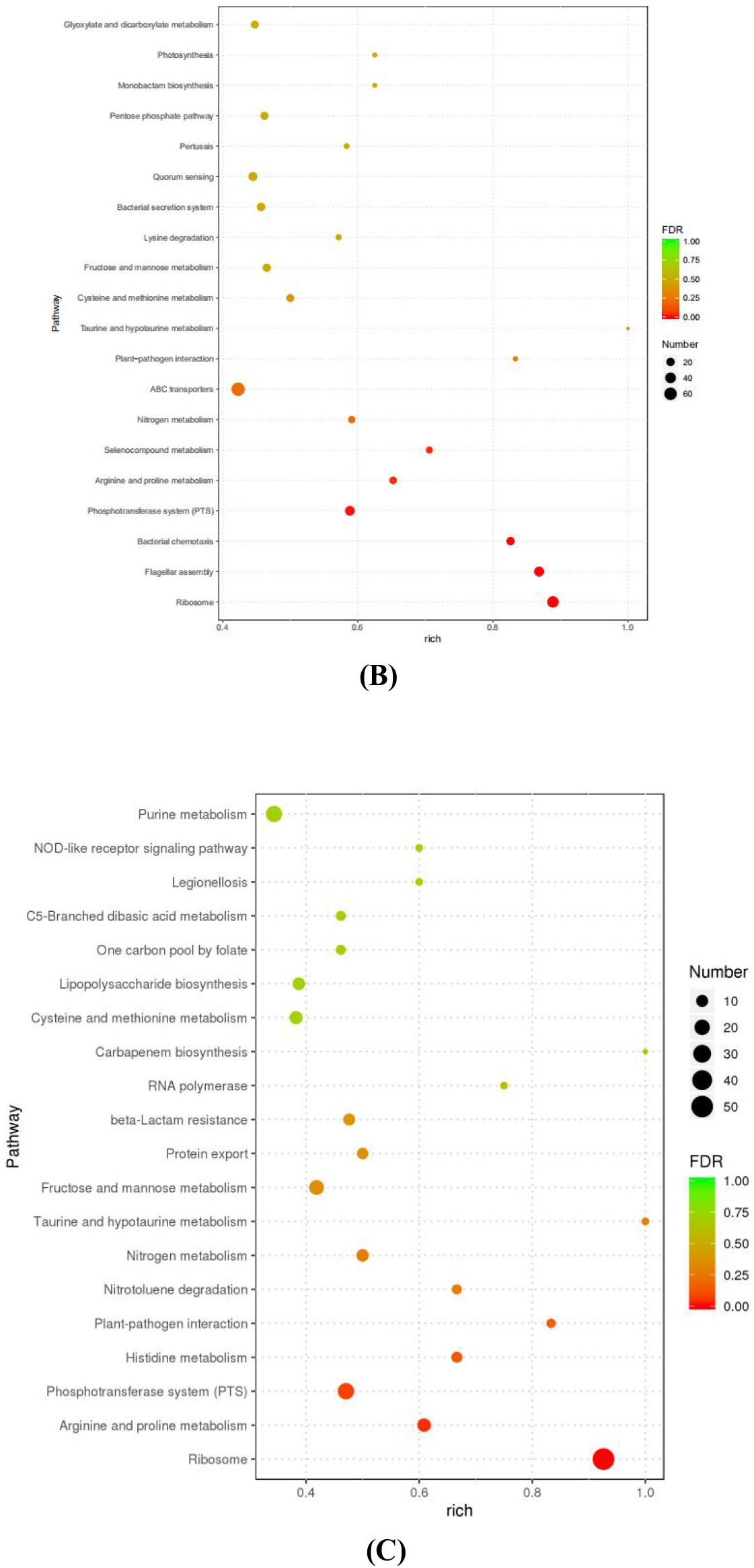
Pathway enrichment scatter plot of DEGs. (A) The top 20 KEGG pathway of B vs C group; (B) The top 20 KEGG pathway of B vs D group; (C) The top 20 KEGG pathway of B vs E group.

**Fig. S4.**
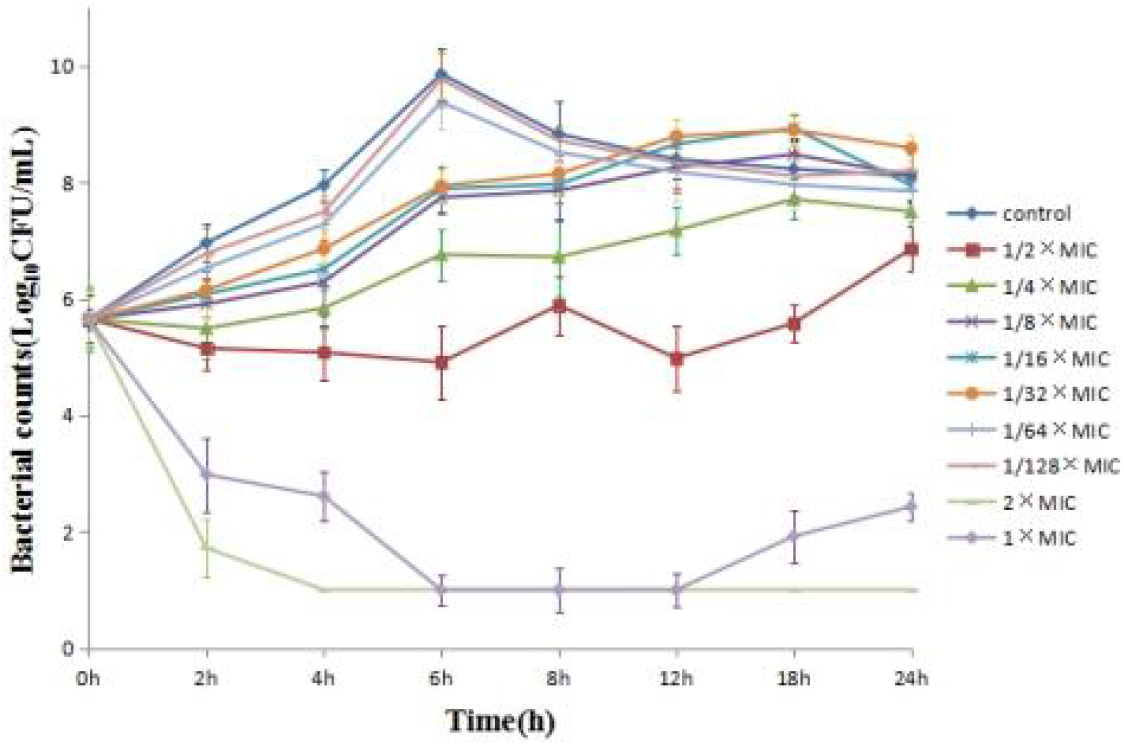
Growth curves of *Salmonella enterica* CICC 21527 in the TSB broth under sub-MIC of enrofloxacin

**Tab. S1.**
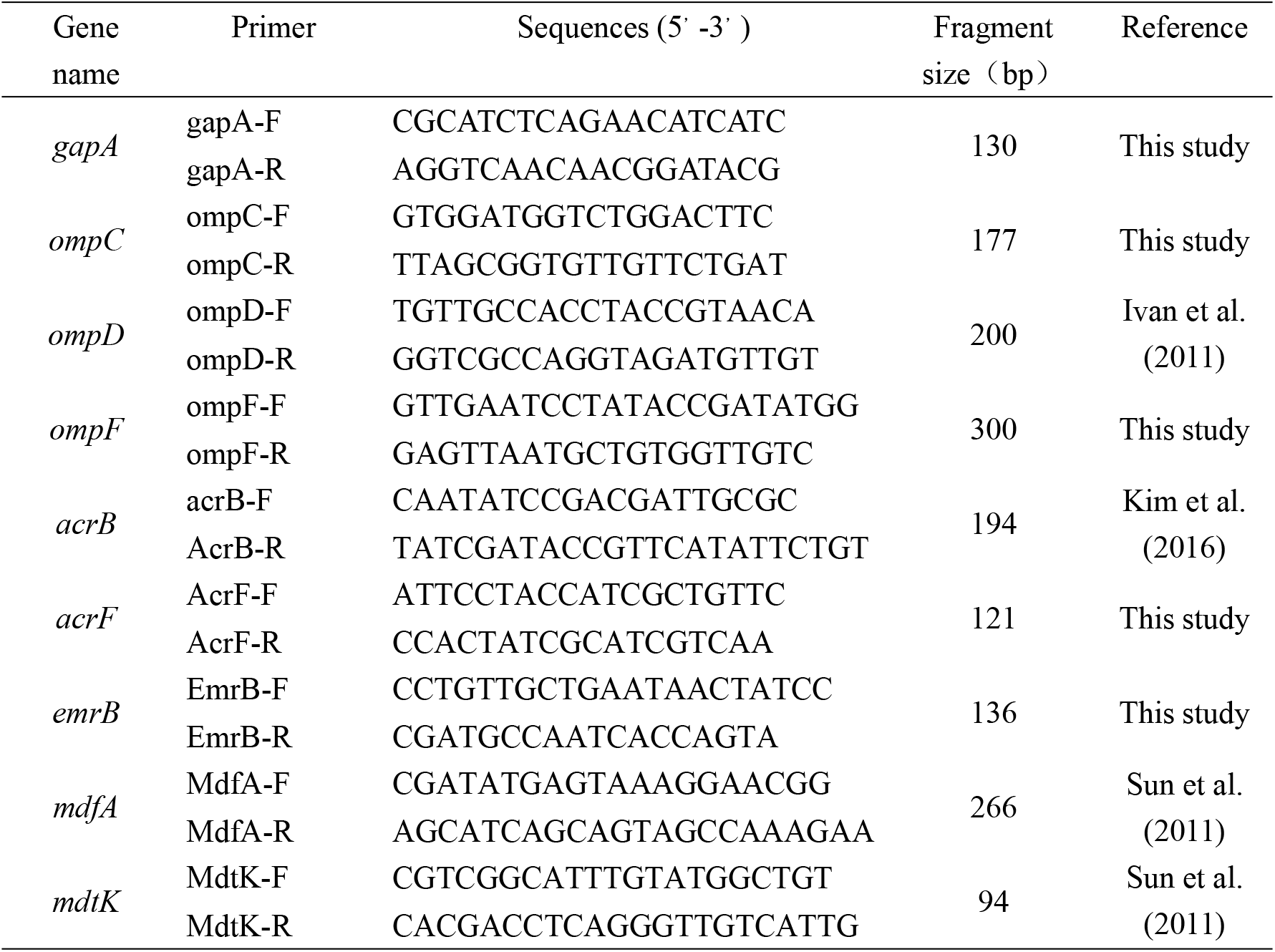
Primers used for identification OMPs and MDR efflux pumps of *Salmonella*.

**Tab. S2.**
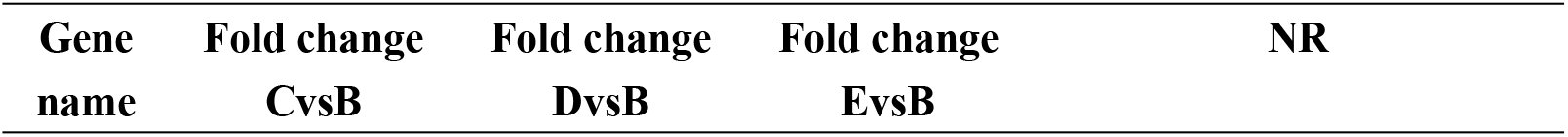
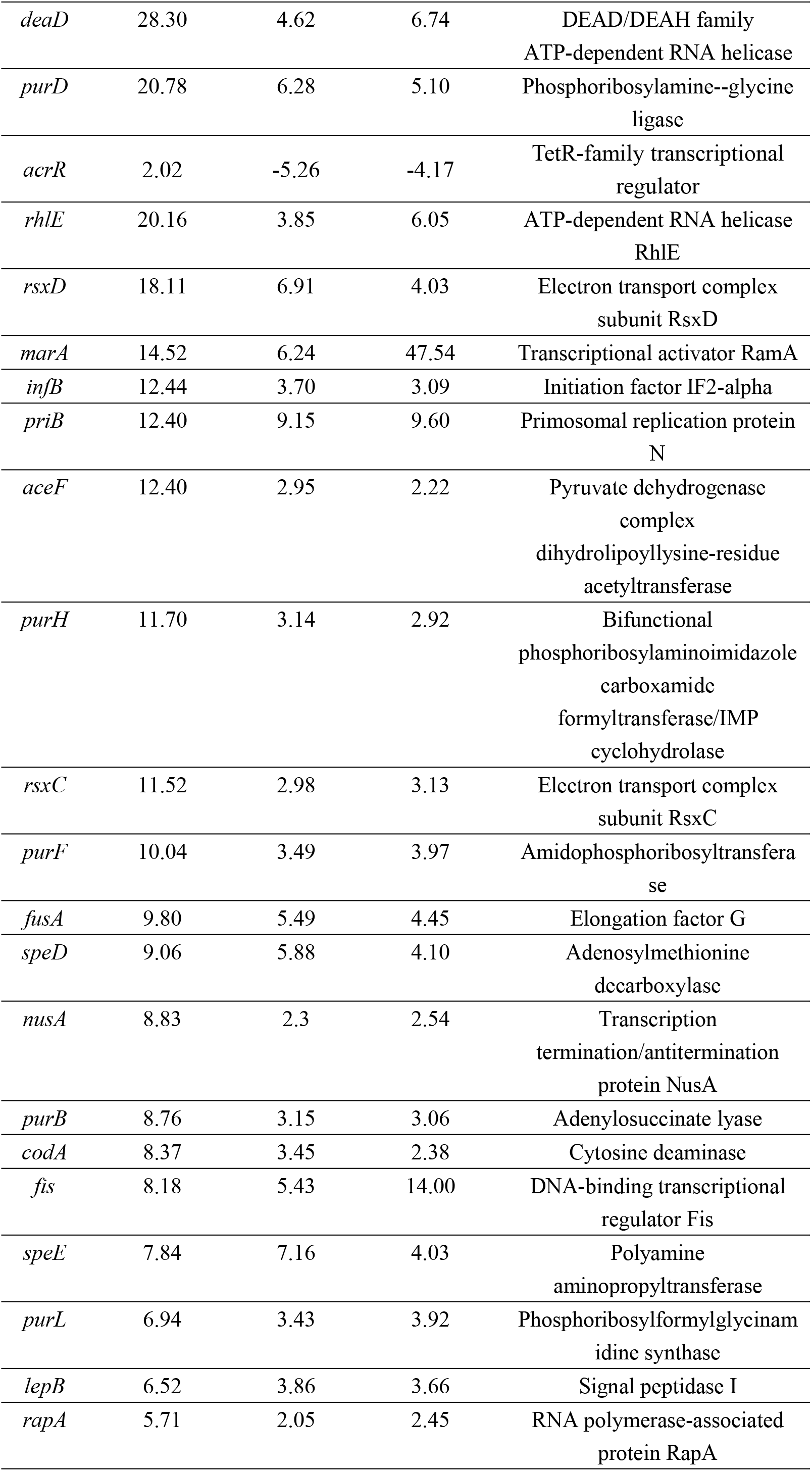
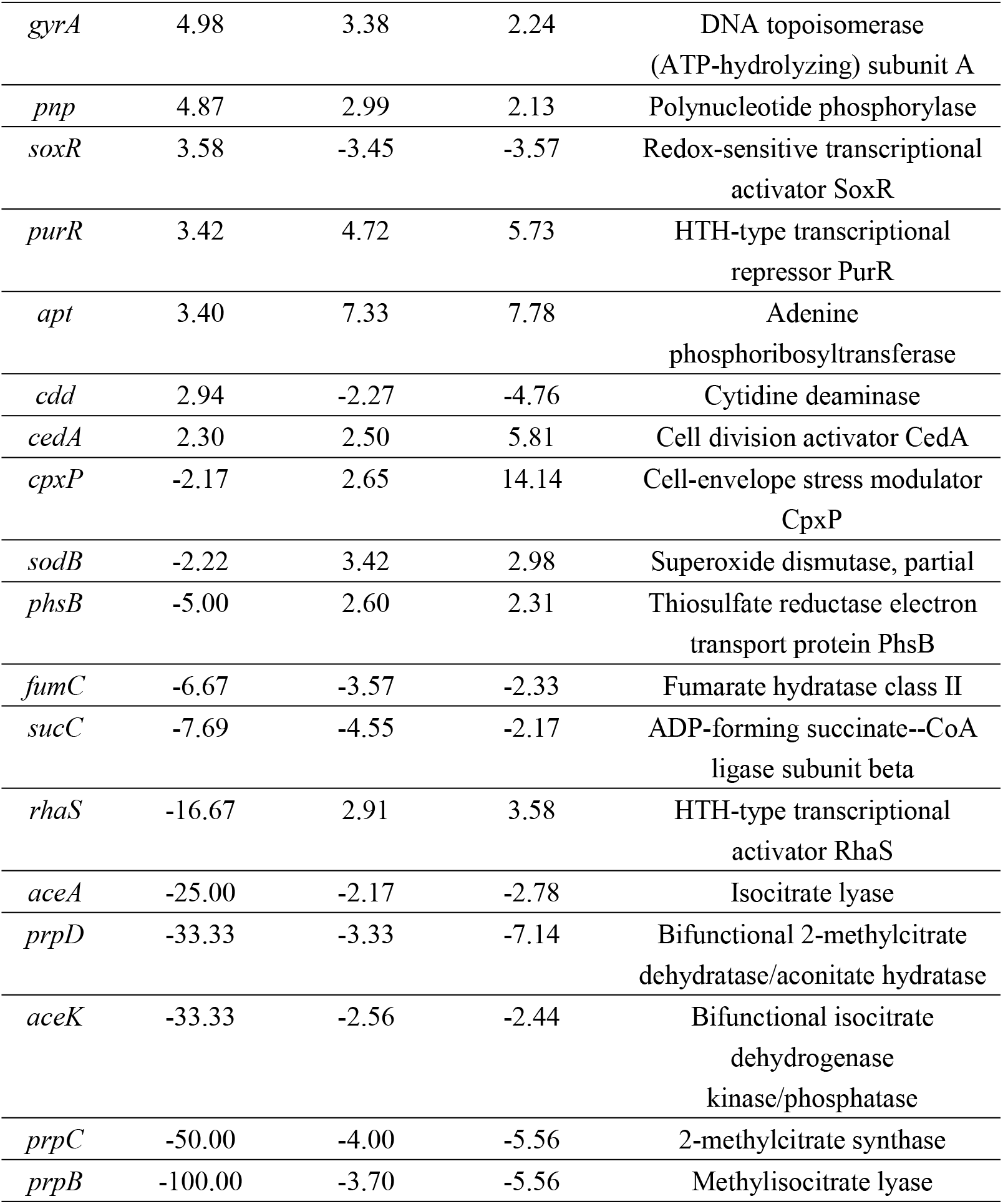
Differently expressed genes between groups of the Co-DEGs among three groups of mutants.

## References

[1] Maria H, Zhao S, James P, et al. Comparative Genomic Analysis and Virulence Differences in Closely Related Salmonella enterica Serotype Heidelberg Isolates from Humans, Retail Meats, and Animals [J]. Genome Biology and Evolution. 2014, (5): 1046–68.

[2] Hendriksen R S, Vieira A R, Karlsmose S, et al. Global monitoring of Salmonella serovar distribution from the World Health Organization Global Foodborne Infections Network Country Data Bank: results of quality assured laboratories from 2001 to 2007 [J]. Foodborne Pathogens and Disease. 2011, 8(8): 887–900.

[3] Martinez M, Mcdermott P, Walker R. Pharmacology of the fluoroquinolones: A perspective for the use in domestic animals [J]. Veterinary Journal. 2006, 172(1): 10–28.

[4] Hopkins K L, Davies R H, Threlfall E J. Mechanisms of quinolone resistance in Escherichia coli and Salmonella: Recent developments [J]. International Journal of Antimicrobial Agent. 2005, 25(5): 358–73.

[5] Lo N W S, Chu M T, Ling J M. Increasing quinolone resistance and multidrug resistant isolates among Salmonella enterica in Hong Kong [J]. Journal of Infection. 2012, 65(6): 528–40.

[6] FáBrega A, Madurga S, Giralt E, et al. Mechanism of action of and resistance to quinolones [J]. Microbial Biotechnology. 2009, 2(1): 40–61.

[7] Rushdy A A, Mabrouk M I, Abu-Sef A H, et al. Contribution of different mechanisms to the resistance to fluoroquinolones in clinical isolates of Salmonella enterica [J]. Brazilian Journal of Infectious Diseases. 2013, 17(4): 431–7.

[8] Nishino K, Latifi T, Groisman E A. Virulence and drug resistance roles of multidrug efflux systems of Salmonella enterica serovar Typhimurium [J]. Molecular Microbiology. 2010, 59(1): 126–41.

[9] Seiji Y, Saya N, Aiko F, et al. Cooperation of the multidrug efflux pump and lipopolysaccharides in the intrinsic antibiotic resistance of Salmonella enterica serovar Typhimurium [J]. Journal of Antimicrobial Chemotherapy. 2013, (5): 1066–70.

[10] Andersson D I, Hughes D. Microbiological effects of sublethal levels of antibiotics [J]. Nature Reviews Microbiology. 2014, 12(7): 465.

[11] Chow L K M, Ghaly T M, Gillings M. A survey of sub-inhibitory concentrations of antibiotics in the environment [J]. Journal of Environmental. 2021, 99: 21–7.

[12] Li L, Dai X, Wang Y, et al. RNA-seq-based analysis of drug-resistant Salmonella enterica serovar Typhimurium selected in vivo and in vitro [J]. Plos One. 2017, 12(4): e0175234.

[13] Horst M, Schuurmans J M, Smid M C, et al. De novo acquisition of resistance to three antibiotics by Escherichia coli [J]. Microbial drug resistance. 2011, 17(2): 141–7.

[14] Thi T D, López E, RodríGuez-Rojas A, et al. Effect of recA inactivation on mutagenesis of Escherichia coli exposed to sublethal concentrations of antimicrobials [J]. Journal of Antimicrobial Chemotherapy. 2011, 66(3): 531.

[15] Kohanski M A, Depristo M A, Collins J J. Sublethal antibiotic treatment leads to multidrug resistance via radical-induced mutagenesis [J]. Molecular cell. 2010, 37(3): 311–20.

[16] Balashov S, Humayun M Z. Mistranslation induced by streptomycin provokes a RecABC/RuvABC-dependent mutator phenotype in Escherichia coli cells [J]. Journal of Molecular Biology. 2002, 315(4): 513–27.

[17] López E, Elez M, Matic I, et al. Antibiotic-mediated recombination: ciprofloxacin stimulates SOS-independent recombination of divergent sequences in Escherichia coli [J]. Molecular Microbiology. 2010, 64(1): 83–93.

[18] Beaber J W, Hochhut B, Waldor M K. SOS response promotes horizontal dissemination of antibiotic resistance genes [J]. Nature. 2004, 427(6969): 72–4.

[19] Wistrand-Yuen E, Knopp M, Hjort K, et al. Evolution of high-level resistance during low-level antibiotic exposure [J]. Nature Communications. 2018, 9(1): 1599.

[20] Zhang C-Z, Ren S-Q, Chang M-X, et al. Resistance mechanisms and fitness of Salmonella Typhimurium and Salmonella Enteritidis mutants evolved under selection with ciprofloxacin in vitro [J]. Scientific reports. 2017, 7(1): 9113.

[21] Dawan J, Uddin M J, Ahn J. Development of de novo resistance in Salmonella Typhimurium treated with antibiotic combinations [J]. FEMS microbiology letters. 2019, 366(10): fnz127.

[22] Wayne P A. CLINICAL AND LABORATORY STANDARDS INSTITUTE. PERFORMANCE STANDARDS FOR ANTIMICROBIAL SUSCEPTIBILITY TESTING [J]. 2011.

[23] Gullberg E, Cao S, Berg O G, et al. Selection of Resistant Bacteria at Very Low Antibiotic Concentrations [J]. PLoS Pathogens. 2011, 7(7): 1–9.

[24] Gu Y, Wang S, Huang L, et al. Development of Resistance in Escherichia coli ATCC25922 under Exposure of Sub-Inhibitory Concentration of Olaquindox[J]. Antibiotics. 2020, 9(11):791.

[25] Samuel H, Albert R, Peter D, et al. Serotypes and Antimicrobial Resistance in Salmonella enterica Recovered from Clinical Samples from Cattle and Swine in Minnesota, 2006 to 2015 [J]. Plos One. 2016, 11(12): e0168016.

[26] Kim S Y, Lee S K, Park M S, et al. Analysis of the Fluoroquinolone Antibiotic Resistance Mechanism of Salmonella enterica Isolates [J]. Journal of Microbiology and Biotechnology. 2016, 26(9): 1605–1612.

[27] Dimitrov T, Dashti A A, Albaksami O, et al. Ciprofloxacin-resistant Salmonella enterica serovar typhi from Kuwait with novel mutations in gyrA and parC genes [J]. Journal of clinical microbiology. 2009, 47(1): 208–11.

[28] O’regan E, Quinn T, Pagès J-M, et al. Multiple regulatory pathways associated with high-level ciprofloxacin and multidrug resistance in Salmonella enterica serovar enteritidis: involvement of RamA and other global regulators [J]. Antimicrobial agents and chemotherapy. 2009, 53(3): 1080–7.

[29] Qian H, Cheng S, Liu G, et al. Discovery of seven novel mutations of gyrB, parC and parE in Salmonella Typhi and Paratyphi strains from Jiangsu Province of China [J]. Scientific Reports. 2020, 10(1): 7359.

[30] Baucheron S, Chaslus-Dancla E, Cloeckaert A, et al. High-Level Resistance to Fluoroquinolones Linked to Mutations in gyrA, parC, and parE in Salmonella enterica Serovar Schwarzengrund Isolates from Humans in Taiwan[J]. Antimicrobial Agents and Chemotherapy. 2005, 49(2):862–863.

[31] Stephen B, Thanh D P, Thieu N, et al. Fitness benefits in fluoroquinolone-resistant Salmonella Typhi in the absence of antimicrobial pressure [J]. eLife. 2013, 2: e01229.

[32] Ching C, Zaman M H. Development and selection of low-level multi-drug resistance over an extended range of sub-inhibitory ciprofloxacin concentrations in Escherichia coli [J]. Scientific Reports. 2020, 10(1): 8754.

[33] Sonia, Morgan-Linnell Lauren, et al. Mechanisms accounting for fluoroquinolone resistance in Escherichia coli clinical isolates [J]. Antimicrobial Agents and Chemotherapy. 2009, 53(1): 235–241.

[34] Kang H-W, Woo G-J. Increase of multidrug efflux pump expression in fluoroquinolone-resistant Salmonella mutants induced by ciprofloxacin selective pressure [J]. Research in veterinary science, 2014, 97(2): 182–6.

[35] Du X, Liu Y, Sun X, et al. The mRNA expression of ompF, invA and invE was associated with the ciprofloxacin-resistance in Salmonella [J]. Archives of Microbiology. 2020, 202, 2263–2268.

[36] Singh P R, Ceccarelli M, Lovelle M, et al. Antibiotic Permeation across the OmpF Channel: Modulation of the Affinity Site in the Presence of Magnesium [J]. The Journal of Physical chemistry. 2012, 116(15): 4433–8.

[37] Cama J, Bajaj H, Pagliara S, et al. Quantification of Fluoroquinolone Uptake through the Outer Membrane Channel OmpF of Escherichia coli [J]. Journal of American Chemical Society 2015, 137(43): 13836–43.

[38] Ferreira M, Sousa C F, Gameiro P. Fluoroquinolone Metalloantibiotics to Bypass Antimicrobial Resistance Mechanisms: Decreased Permeation through Porins [J]. Membranes. 2020, 11(1): 3.

[39] Tsukasa H, Akihito Y, Kunihiko N. TolC dependency of multidrug efflux systems in Salmonella enterica serovar Typhimurium[J]. Journal of Antimicrobial Chemotherapy. 2010(7):1372–1376.

[40] Singh R, Swick M C, Ledesma K R, et al. Temporal Interplay between Efflux Pumps and Target Mutations in Development of Antibiotic Resistance in Escherichia coli [J]. Antimicrob Agents Chemother. 2012, 56(4):1680–1685.

[41] Chang T M, Lu P L, Li H H, et al. Characterization of Fluoroquinolone Resistance Mechanisms and Their Correlation with the Degree of Resistance to Clinically Used Fluoroquinolones among Escherichia coli Isolates [J]. Journal of Chemotherapy. 2007, 19(5): 488–94.

[42] Zhang C Z, Chen P X, Yang L, et al. Coordinated Expression of acrAB-tolC and Eight Other Functional Efflux Pumps Through Activating ramA and marA in Salmonella enterica serovar Typhimurium [J]. Microbial Drug Resistance. 2017, 24(2): 120–125.

[43] Jessica M A B, Helen E S, Vito R, et al. Expression of homologous RND efflux pump genes is dependent upon AcrB expression: implications for efflux and virulence inhibitor design [J]. Journal of Antimicrobial Chemotherapy. 2014, (2): 424–31.

[44] Yamasaki S, Nikaido E, Nakashima R, et al. The crystal structure of multidrug-resistance regulator RamR with multiple drugs [J]. Nature Communications. 2013, 4:2078.

[45] Switzer R L, Zalkin H, Saxild H H. Purine, Pyrimidine, and Pyridine Nucleotide Metabolism[M]. John Wiley & Sons, Ltd, 2014.

[46] Breton Y L, Mistry P, Valdes K M, et al. Genome-wide identification of genes required for fitness of group A Streptococcus in human blood.[J]. Infection & Immunity, 2013, 81(3):862–875.

[47] Kim J K, Jang H A, Won Y J, et al. Purine biosynthesis-deficient Burkholderia mutants are incapable of symbiotic accommodation in the stinkbug[J]. Isme Journal, 2014, 8(3):552.

[48] GAO-FEI, YUN-DAN, ZHENG, et al. Novel Mechanistic Insights into Bacterial Fluoroquinolone Resistance [J]. Journal of proteome research. 2019, 18(11): 3955–66.

